# Ultra-fast insulin-pramlintide co-formulation for improved glucose management in diabetic rats

**DOI:** 10.1101/2021.04.12.439573

**Authors:** Caitlin L. Maikawa, Peyton C. Chen, Eric T. Vuong, Leslee T. Nguyen, Joseph L. Mann, Andrea I. d’Aquino, Rayhan A. Lal, David M. Maahs, Bruce A. Buckingham, Eric A. Appel

**Affiliations:** Department of Bioengineering, Stanford University, Stanford CA 94305, USA; Department of Biochemistry, Stanford University, Stanford CA 94305, USA; Department of Materials Science & Engineering, Stanford University, Stanford CA 94305, USA; Department of Medicine (Endocrinology), Stanford University, Stanford CA 94305, USA; Department of Pediatrics (Endocrinology), Stanford University, Stanford CA 94305, USA; Diabetes Research Center, Stanford University, Stanford CA 94305, USA; ChEM-H Institute, Stanford University, Stanford CA 94305, USA

**Keywords:** insulin, amylin, hormones, diabetes, drug delivery

## Abstract

Dual-hormone replacement therapy with insulin and amylin in patients with type 1 diabetes has the potential to improve glucose management. Unfortunately, currently available formulations require burdensome separate injections at mealtimes and have disparate pharmacokinetics that do not mimic endogenous co-secretion. Here, we use amphiphilic acrylamide copolymers to create a stable co-formulation of monomeric insulin and amylin analogues (lispro and pramlintide) with synchronous pharmacokinetics and ultra-rapid action. The co-formulation is stable for over 16 hours under stressed aging conditions, whereas commercial insulin lispro (Humalog) aggregates in 8 hours. The faster pharmacokinetics of monomeric insulin in this co-formulation resulted in increased insulin-pramlintide overlap of 75 ± 6% compared to only 47 ± 7% for separate injections. The co-formulation resulted in similar delay in gastric emptying compared to pramlintide delivered separately. In a glucose challenge, in rats the co-formulation reduced deviation from baseline glucose compared to insulin only, or separate insulin and pramlintide administrations. Further, comparison of interspecies pharmacokinetics of monomeric pramlintide suggests that pharmacokinetics observed for the co-formulation will be well preserved in future translation to humans. Together these results suggest that the co-formulation has the potential to improve mealtime glucose management and reduce patient burden in the treatment of diabetes.

## 1. Introduction

Patients with type 1 diabetes lack the ability to produce both endogenous insulin and amylin after an autoimmune response destroys the pancreatic beta-cells.[1] In individuals without diabetes, insulin and amylin work synergistically to control post-prandial glucose; amylin delays gastric emptying and suppresses glucagon action, while insulin promotes cellular glucose uptake.[1–2] Studies have shown that dual-hormone replacement therapy with insulin and amylin results in improved glycemic outcomes for individuals with diabetes, including a 0.3% reduction in HbA1c compared to treatment with insulin alone.[3] However, treatment of type 1 diabetes over the last 100 years has primarily focused on insulin replacement. While a commercially available amylin analogue (pramlintide) exists, only 1.5% of patients who would benefit from amylin replacement therapy had adopted it by 2012.[4] This is primarily due to formulation challenges that result in the need for a burdensome separate injection of amylin in addition to insulin at mealtimes.

Amylin is highly unstable and rapidly aggregates to form inactive and immunogenic amyloid fibrils.[5] Pramlintide, the only commercially available amylin analogue, has three amino acid modifications to reduce its propensity to aggregate into amyloid fibrils, thus improving its shelf-life. Unfortunately, pramlintide is typically formulated at pH 4, making it incompatible with current rapid-acting insulin formulations (pH~7).[2] Further, in current clinical administrations insulin and pramlintide have disparate pharmacokinetics, which is in contrast to endogenous cosecretion of the two hormones from the beta-cells following the same diurnal patterns.[6] We hypothesize this difference in kinetics results in reduced synergistic effects and requires pramlintide doses greater than physiological ratios of insulin to pramlintide. The difference in absorption kinetics when delivered exogenously results from the different association states of insulin and pramlintide in formulation (Figure 1a). Pramlintide only exists as a monomer, while insulin formulations contain a mixture of hexamers, dimers, and monomers.[7] The mixture of insulin association states, specifically the presence of the insulin hexamer that is responsible for the subcutaneous insulin depot effect, results in delayed absorption and prolonged duration of insulin action.[7]

**Figure 1.**
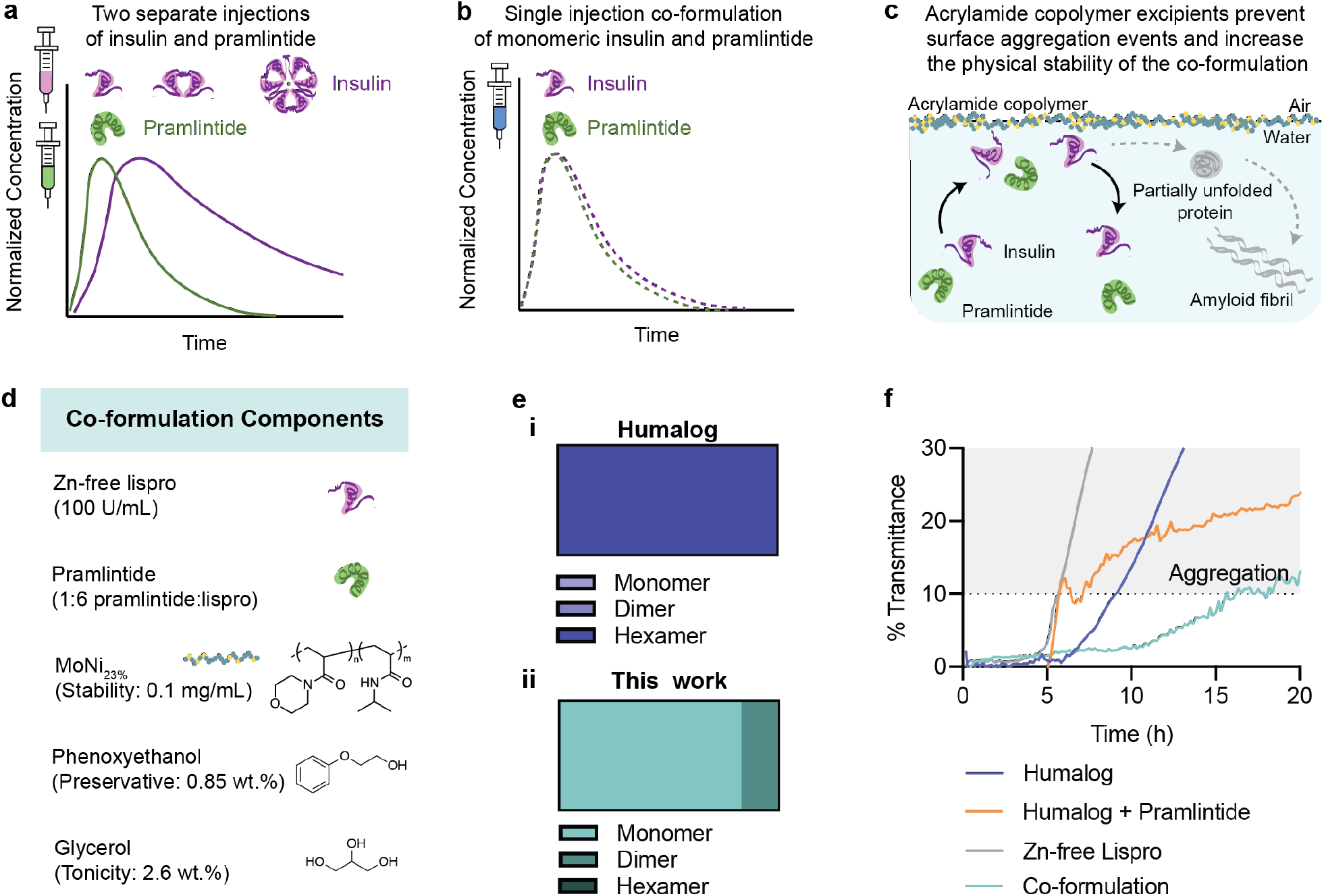
Scheme of formulation kinetics and stability. **a,** Current dual-hormone replacement of insulin and pramlintide requires two separate injections at mealtimes (idealized data for illustration based on reported pharmacokinetics[9]). Not only is this additional injection burdensome, but there is a kinetic mismatch between insulin and pramlintide when delivered exogenously compared to endogenous co-secretion from the beta-cells. This results from the mixed insulin association states present in rapid-acting insulin formulations where monomers and dimers are rapidly absorbed, but the slow dissociation of the insulin hexamer causes extended duration of action. **b,** A single injection co-formulation of monomeric insulin and pramlintide would reduce patient burden, and have better pharmacokinetic overlap that more closely mimics endogenous secretion from the healthy pancreas (idealized data for illustration of study goals). **c,** Amphiphilic acrylamide copolymer excipients can be used to stabilize an insulin-pramlintide co-formulation. These excipients preferentially adsorb onto the air-water interface, displacing insulin and/or pramlintide and preventing the nucleation of aggregation events that initiate amyloid fibril formation. **d,** Co-formulation components. **e,** Insulin association states in (i) Humalog (adapted from the literature^4^) compared to (ii) zinc-free lispro with phenoxyethanol (0.85 wt.%) and glycerol (2.6 wt.%). **f,** Formulation stability in a stressed aging assay (continuous agitation, 37 °C) of (i) Humalog, (ii) Humalog + pramlintide (1:6 pramlintide:lispro), (iii) zinc-free lispro (100U/mL lispro, 0.85 wt.% phenoxyethanol, 2.6 wt.% glycerol, 0.1 mg/mL MoNi_23%_), (iv) Co-formulation (100 U/mL lispro, 1:6 pramlintide:lispro, 0.85 wt.% phenoxyethanol, 2.6 wt.% glycerol, 0.1 mg/mL MoNi23%). Change in transmittance is shown from baseline transmittance. Aggregation is defined as a change in transmittance >10%.

Recent work from our group has exploited non-covalent PEGylation to create an insulin-pramlintide co-formulation where supramolecular modification of both proteins simultaneously with a designer excipient cucurbit[7]uril-poly(ethylene glycol) (CB[7]-PEG) enables stable co-formulation of insulin and pramlintide for delivery in a single administration.[8] This formulation showed increased pharmacokinetic overlap in diabetic pigs, where pramlintide action is slightly extended by formulation with CB[7]-PEG to more closely match subcutaneous insulin absorption.[8] These more similar pharmacokinetics resulted in improved glucagon suppression in diabetic pigs;[8] however, the increased pharmacokinetic overlap was achieved primarily by delaying pramlintide absorption - slowing the pramlintide pharmacokinetic profile – to better overlap the insulin and pramlintide exposure curves. Ideally, a meal-time insulin-pramlintide co-formulation would have ultrafast kinetics of both insulin and pramlintide, allowing both rapid onset and reduced duration of action for both therapeutic proteins (Figure 1b). An insulin drug product with these characteristics would allow for rapid management of meal-time glucose spikes and reduced risk of post-prandial hypoglycemia. In combination with a real-time continuous glucose sensor, this insulin-pramlintide co-formulation with more “on-off” kinetics would provide a significant benefit to automated insulin delivery (“artificial pancreas” systems).

Since our initial non-covalent PEGylation studies, our group has developed amphiphilic acrylamide carrier-dopant copolymer (AC/DC) excipients that are composed of a water soluble “carrier” monomer and a hydrophobic “dopant” monomer.[10] These copolymer excipients prevent protein aggregation at hydrophobic interfaces, such as the air-water interface, and have been used to enable a stable monomeric insulin formulation that exhibited ultrafast insulin pharmacokinetics in diabetic pigs.[10] Typically, insulin aggregation is initiated at the air-water interface by interactions between partially unfolded insulins adsorbed to the interface.[11] These novel amphiphilic acrylamide copolymers preferentially adsorb to the air-water interface, displacing insulin and preventing the nucleation of insulin aggregation events (Figure 1c).[12] These copolymer excipients are advantageous over approaches to non-covalent PEGylation because they lack specific protein-polymer interactions, imbuing stability without altering protein pharmacokinetics. Here, we develop an ultra-fast insulin-amylin co-formulation by leveraging a top-performing acrylamide copolymer excipient acryloylmorpholine-co-N-isopropylacrylamide (MoNi_23%_) to stabilize the two hormones together in formulation. We hypothesize that combining monomeric insulin and pramlintide will result in an ultra-fast insulin pharmacokinetic profile that will better overlap with pramlintide pharmacokinetics to better mimic endogenous co-secretion of the two hormones (Figure 1b). Further, we anticipate the addition of MoNi_23%_, will imbue stability and allow these two hormones to coexist in a single formulation exhibited enhanced stability when compared with commercial insulin drugs.

## 2. Results

### 2.1. Stabilization of an insulin-pramlintide co-formulation

Our previous work has demonstrated the utility of MoNi_23%_ as a stabilizing excipient for monomeric insulin.[10] The propensity of insulin and pramlintide to aggregate to form amyloid fibrils, which are primarily initiated at hydrophobic interfaces, makes them strong candidates for stabilization using MoNi_23%_. It has been shown that MoNi_23%_ can disrupt insulin-insulin interactions at the air-water interface. We hypothesized that we could use MoNi_23%_ to physically stabilize an ultrafast mealtime insulin-pramlintide co-formulation. This co-formulation will use the excipients previously identified in our ultrafast absorbing insulin lispro formulation to promote the insulin monomer, combined with pramlintide to enable a single formulation with increased pharmacokinetic overlap between these two hormones.

As previously reported, zinc-free lispro in the presence of glycerol (2.6 wt.%) and phenoxyethanol (0.85 wt.%) as tonicity and antimicrobial agents, results in a formulation with a high monomer content.[13] Using size-exclusion chromatography with multi-angle light scattering (SEC-MALS) we observed 83% monomers, 17% dimers and 0% hexamers in formulation (Figure 1e, Figure S1). In comparison, commercial Humalog is >99% hexameric.[10] For SEC-MALS measurements, insulin association state is tested alone with only small molecule excipients because both pramlintide and the MoNi_23%_ excipient are of similar molecular weight and would prevent the calculation of monomer content in formulation. The addition of MoNi_23%_ has been shown not to alter the insulin association state by diffusion-ordered nuclear magnetic resonance spectroscopy (DOSY-NMR).[10] Based on our previous results, it is not anticipated that the presence of pramlintide would alter the insulin association state.[8]

The insulin-pramlintide co-formulation is composed of zinc-free lispro (100 U/mL), pramlintide (1:6 molar ratio pramlintide:lispro), glycerol (2.6 wt.%), phenoxyethanol (0.85 wt.%), and MoNi_23%_ (0.1 mg/mL) in phosphate buffer at pH~7 (Figure 1d). A pramlintide ratio of 1:6 was chosen to be consistent with previous work using the CB[7]-PEG stabilized insulin-pramlintide co-formulation in diabetic pigs.[8] Further, a ratio of 1:6 is similar to high endogenous insulin-pramlintide ratios reported in the literature as well as within the range of ratios indicated to be most effective by in silico experiments.[14] Formulation stability was assessed using a stressed aging assay.[8, 10, 12–13, 15] As insulin and/or pramlintide aggregates form, they scatter light which can be measured by absorbance. Here, formulation aggregation is defined as a 10% or greater change in transmittance. Our co-formulation is stable for 16.2 ± 0.1 hours, twice as long as commercial Humalog which aggregates after 8.2 ± 0.5 hours (Figure 1f). The direct addition of pramlintide to Humalog results in a translucent formulation immediately upon mixing which has 5-25% reduced transmittance compared to Humalog alone (Figure S2). This mixture reaches the aggregation threshold after 8 ± 3 hours, which is highly variable due to the variable initial transmittance. Zinc-free lispro alone is mostly monomeric and is highly unstable, aggregating rapidly after 5.7 ± 0.1 hours.

### 2.2 Pharmacokinetics and pharmacodynamics in diabetic rats

After establishing the stability of our insulin-pramlintide co-formulation, we evaluated the pharmacokinetics *in vivo* to determine if the use of monomeric insulin resulted in increased pharmacokinetic overlap. The co-formulation was tested against controls of Humalog alone and separate injections of insulin and pramlintide (Figure 2). A high dose of each formulation (2 U/kg) was given to each rat followed by oral gavage with glucose solution (1 g/kg). A similar magnitude of glucose lowering was observed in all three formulations (Figure 2b). Normalized pharmacokinetic values allow for easier visual comparison of metrics of onset (time to 50% peak up) and duration of action (time to 50% peak down) between formulations (Figure 3a,j). For insulin lispro pharmacokinetics, no statistically significant difference was observed for comparisons of time to onset (time to 50% peak up) or time to peak between formulations (Figure 3a-c). There was a difference in duration of action, defined as 50% of peak down, between formulations (F_2,20_=7.07, P=0.0048). The co-formulation had shorter duration of action (22 ± 2 minutes) compared to separate injections (34 ± 3 minutes, P=0.0034) (Figure 3a,d). Faster onset was also corroborated using exposure ratios - the fraction of the area under the curve (AUC) at a given timepoint over the total (AUC_t_/AUC_120_). The co-formulation showed a greater fraction of total exposure compared to Humalog and separate injections at 6-, 15- and 30-minute timepoints (Figure 3e-i). There was no difference in insulin lispro (F_2,20_=0.53, P=0.59) or pramlintide (F_2,10_=3.27, P=0.10) area under the exposure curve between formulations (Figure S3-4). As expected, there were no differences observed between pramlintide kinetics delivered as separate injections versus in the co-formulation (Figure 3j-m, Figure S4). The shift of the co-formulation insulin lispro pharmacokinetic curve to the left was confirmed by overlaying the insulin pramlintide curves for delivery by separate injections or co-formulation and comparing overlap time (Figure 4). Overlap was defined as the ratio of overlap over total peak width at half peak height (overlap ÷ (lispro + pramlintide − overlap). As hypothesized, delivery of monomeric insulin with pramlintide in a co-formulation resulted in increased overlap (0.75 ± 0.06) compared to separate injections (0.47 ± 0.07, F_1,10_=6.96, P=0.025) (Figure 4c). The faster insulin kinetics and increased overlap between insulin and pramlintide observed in our co-formulation more closely mimic insulin-pramlintide secretion at mealtimes. Further, unlike the dissociation of the insulin hexamer, which shows more rapid dissociation in rodents and pigs compared to humans, the absorption kinetics of the insulin monomer is better preserved when compared between species (Figure S5-S7). This suggests that the ultrafast kinetics, and increased insulin-pramlintide overlap, observed in these studies will translate well to human patients.

**Figure 2.**
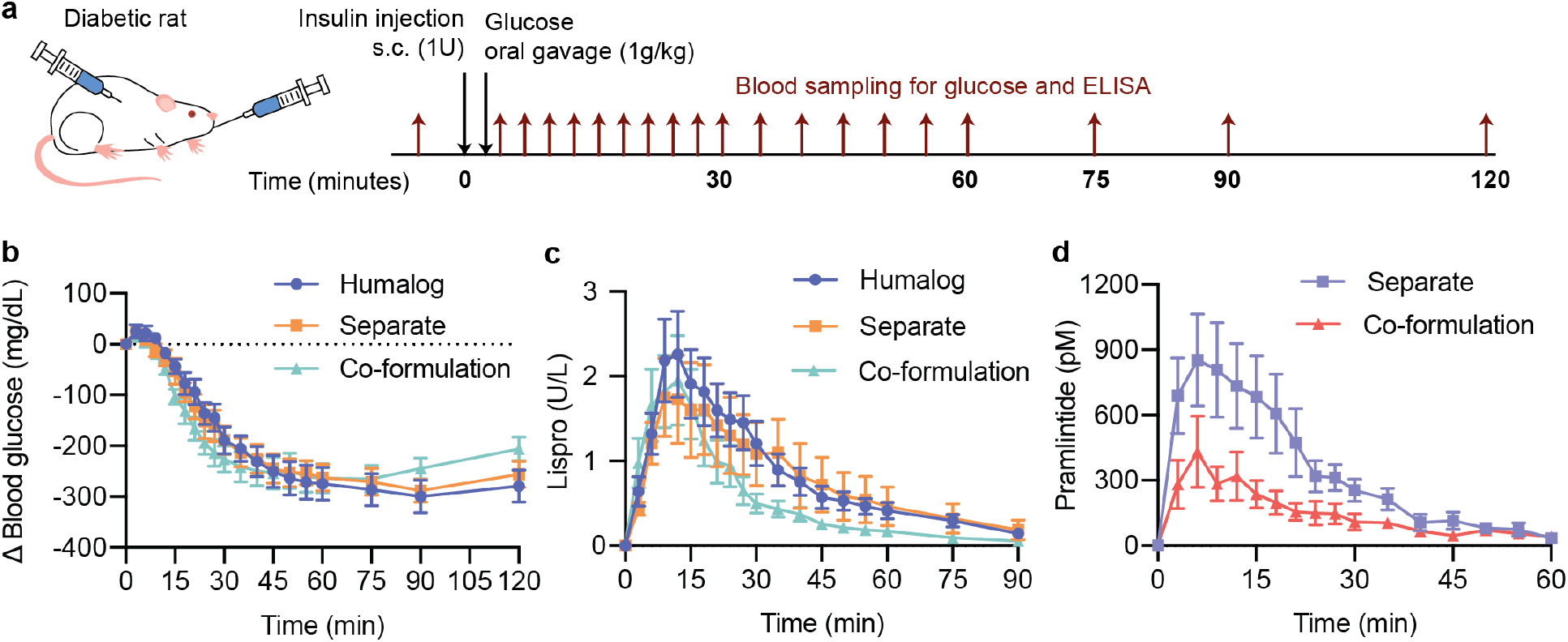
Pharmacokinetics and pharmacodynamics in diabetic rats. Fasted male diabetic rats (n=11) received subcutaneous administration of (i) Humalog, (ii) separate injections of Humalog and pramlintide, or (iii) insulin-pramlintide co-formulation. **a,** Insulin administration was immediately followed with oral gavage with a glucose solution (1 g/kg). Each rat received all treatment groups. **b,** Change in blood glucose levels from baseline following treatment. **c,d,** Pharmacokinetics of (**c)** insulin lispro or **(d)** pramlintide. See Figure S3 and S4 for area under the curve (AUC) exposure comparison for lispro (F_2,20_=0.53, P=0.59) and pramlintide (F_2,10_=3.27, P=0.10).

**Figure 3.**
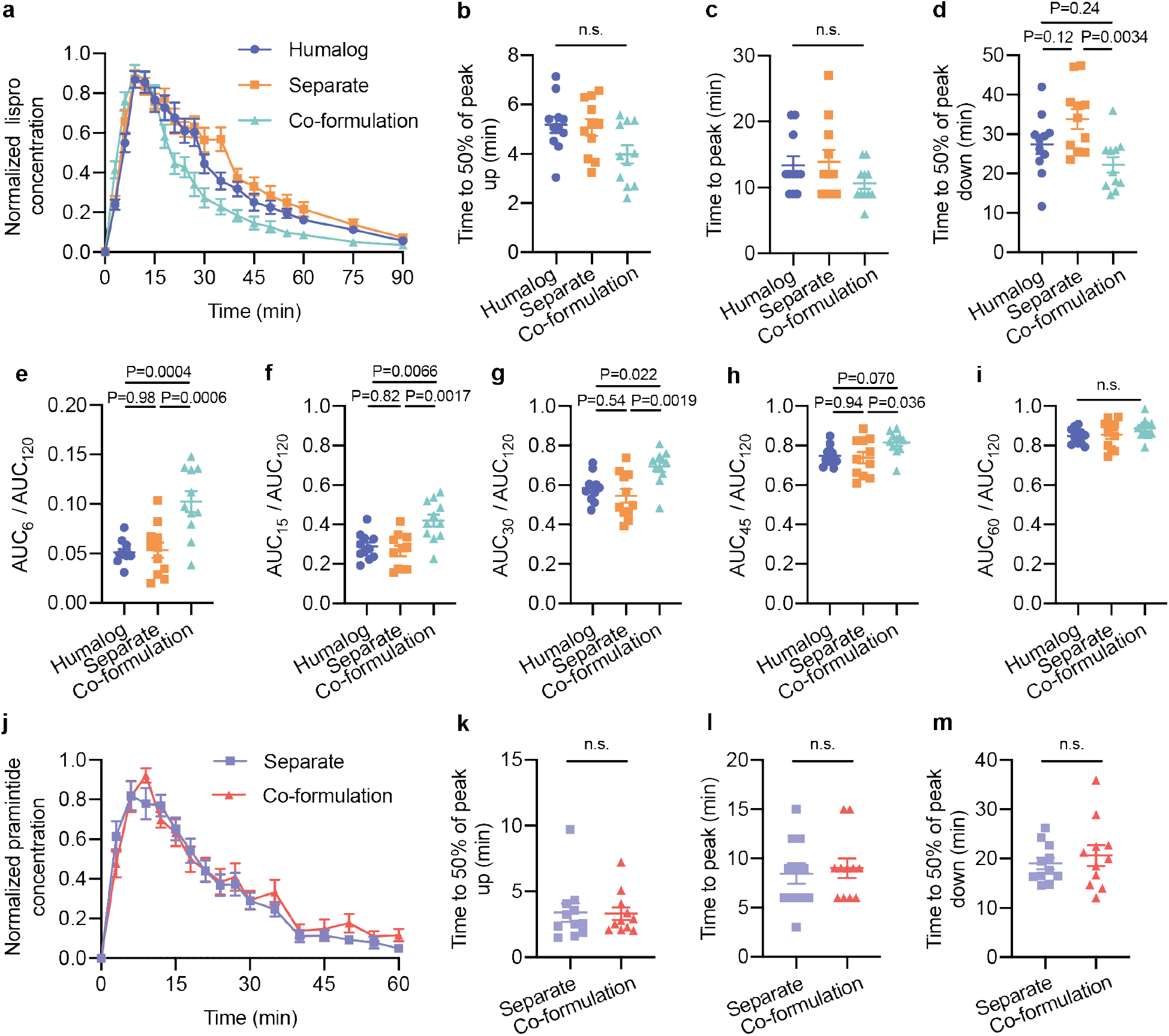
Onset and duration of action in diabetic rats. Fasted male diabetic rats (n=11) received subcutaneous administration of (i) Humalog, (ii) separate injections of Humalog and pramlintide, or (iii) insulin-pramlintide co-formulation. Insulin administration was immediately followed with oral gavage with a glucose solution (1 g/kg). Each rat received all treatment groups. **a,j,** Pharmacokinetics for each rat was individually normalized to the peak serum levels and the normalized values were averaged for **(a)** insulin lispro or **(j)** pramlintide. **b,k,** Exposure onset defined as time to 50% of the peak up for **(b)** insulin lispro or **(k)** pramlintide. **c,l,** Exposure peak for **(c)** insulin lispro or **(l)** pramlintide. **d,m,** Exposure onset defined as time to 50% of the peak up for **(d)** insulin lispro or **(m)** pramlintide. **e-i,** Fraction of lispro exposure as a ratio of AUC_t_/AUC_120_ at **e,** t=6; **f,** t=15; **g,** t=30; **h,** t=45; **i,** t=60. Statistical significance was determined by restricted maximum likelihood repeated measures mixed model. Tukey HSD post-hoc tests were applied to account for multiple comparisons (b-i, k-m). Bonferroni post hoc tests were performed to account for comparisons of multiple individual exposure time points, and significance and a were adjusted (α= 0.01) (e-i).

**Figure 4.**
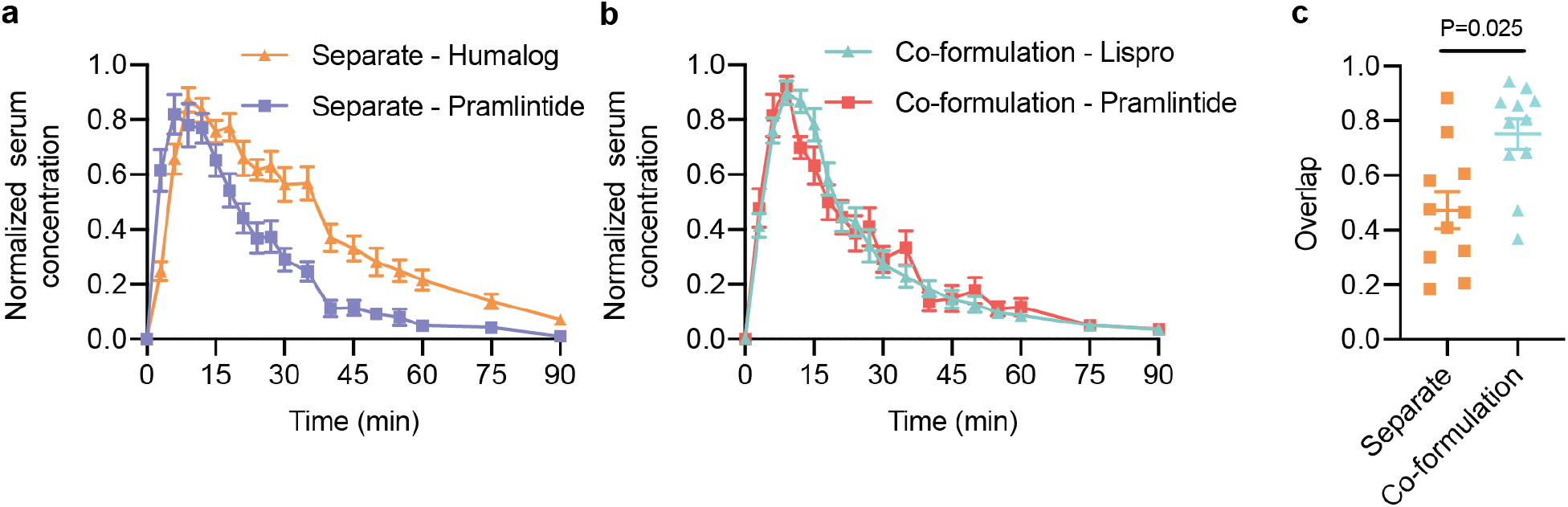
Pharmacokinetic overlap of formulations. **a,b,** Average normalized serum concentrations (for each rat, n=11/group) for insulin and pramlintide when delivered **(a)** as two separate injections and **(b)** when delivered together as a co-formulation. **c,** Overlap between the two curves was defined as the total time spent above 0.5 for both insulin and pramlintide curves (width at half-peak height), shown as a ratio of the overlap time to the total width of both peaks (overlap ÷ (lispro + pramlintide − overlap). Statistical significance was determined by restricted maximum likelihood repeated measures mixed model.

### 2.3 Gastric emptying of acetaminophen in diabetic rats

With our co-formulation in hand, we sought to determine if there were mealtime benefits to our co-formulation compared to standard administrations of Humalog alone or Humalog and pramlintide administered separately. First, we used acetaminophen as model cargo to confirm pramlintide function by testing its ability to delay gastric emptying after formulation administration (Figure 5). We expected that pramlintide in both separate administrations and in the co-formulation would result in delayed gastric emptying compared to Humalog alone. Indeed, the time to peak acetaminophen concentration was delayed until 76 ± 5 minutes for separate injections and 68 ± 6 minutes for the co-formulation compared to 35 ± 5 minutes for Humalog alone, demonstrating there was no difference in time to peak acetaminophen between separate injections and the co-formulation (Figure 5c).

**Figure 5.**
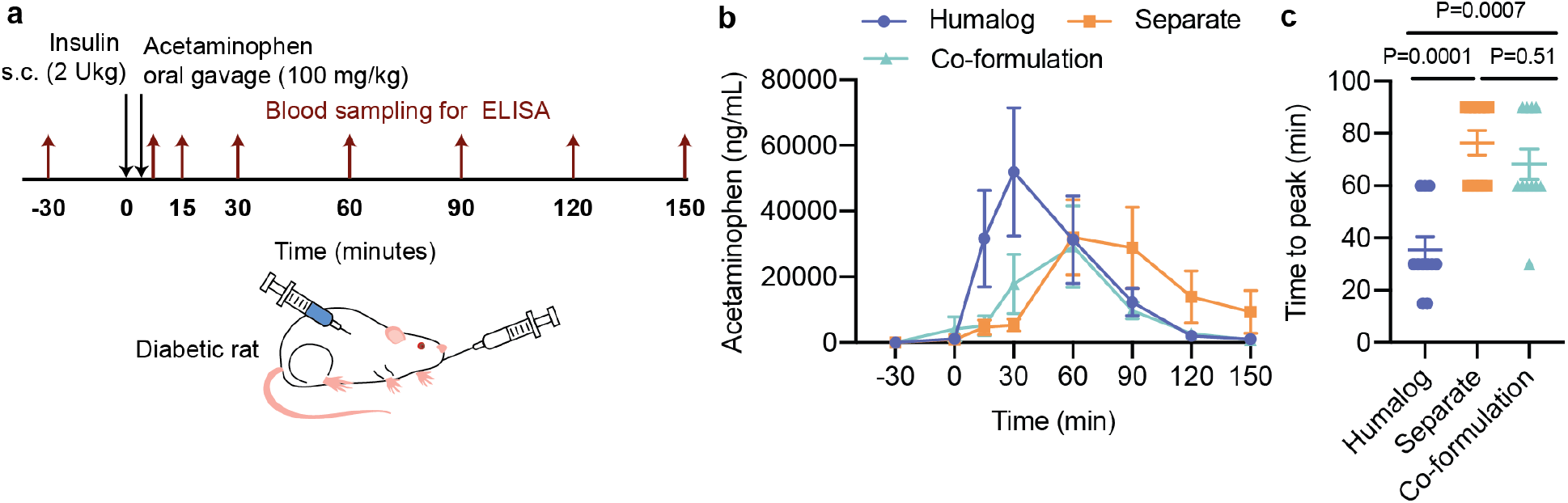
Gastric emptying in diabetic rats. Fasted male diabetic rats received subcutaneous administration of (i) Humalog, (ii) separate injections of Humalog and pramlintide, or (iii) insulin-pramlintide co-formulation. **a,** Gastric emptying experiment where insulin administration (2 U/kg) was immediately followed with oral gavage with an acetaminophen slurry (100 mg/kg). Each rat (n=11) received all treatment groups. **b,** Acetaminophen serum concentration. **c,** Time to peak exposure of acetaminophen serum concentration. All data is shown as mean ± SE. Statistical significance was determined by restricted maximum likelihood repeated measures mixed model. Tukey HSD post-hoc tests were applied to account for multiple comparisons.

### 2.4 Mealtime glucose challenge in diabetic rats

We further tested the co-formulation in a simulated mealtime challenge with a low dose of subcutaneous insulin (0.75 U/kg) and a high dose of glucose (2 g/kg) administered by oral gavage (Figure 6). Starting glucose was variable between rats but was similar for each of the three formulations within a rat (See Figure S8-9 for individual glucose curves). In contrast to the glucose measurements in the pharmacokinetic experiments where insulin was dominant, this experiment aimed to reduce the insulin dose and increase the glucose load to better simulate mealtime glucose management. Yet, the insulin-to-carbohydrate ratio dosed here was still not ideally matched due to the practical constraints of accurately administering small volumes of insulin to the rats. All three formulations had similar control of the glucose peak (Figure 6c). When looking specifically at the co-formulation, we observe control of this mealtime glucose spike while also reducing the magnitude of glucose lowering below baseline levels (Figure 6b,d). In contrast, while the delayed gastric emptying for the separate injection formulations results in rapid lowering of glucose levels and control of the mealtime glucose spike, it also results in a greater glucose drop below baseline. The Humalog-only administration results in a similar glucose curve to separate administrations of insulin and pramlintide but with delayed glucose lowering since glucose release is not slowed as in the other formulations on account of the pramlintide.

**Figure 6.**
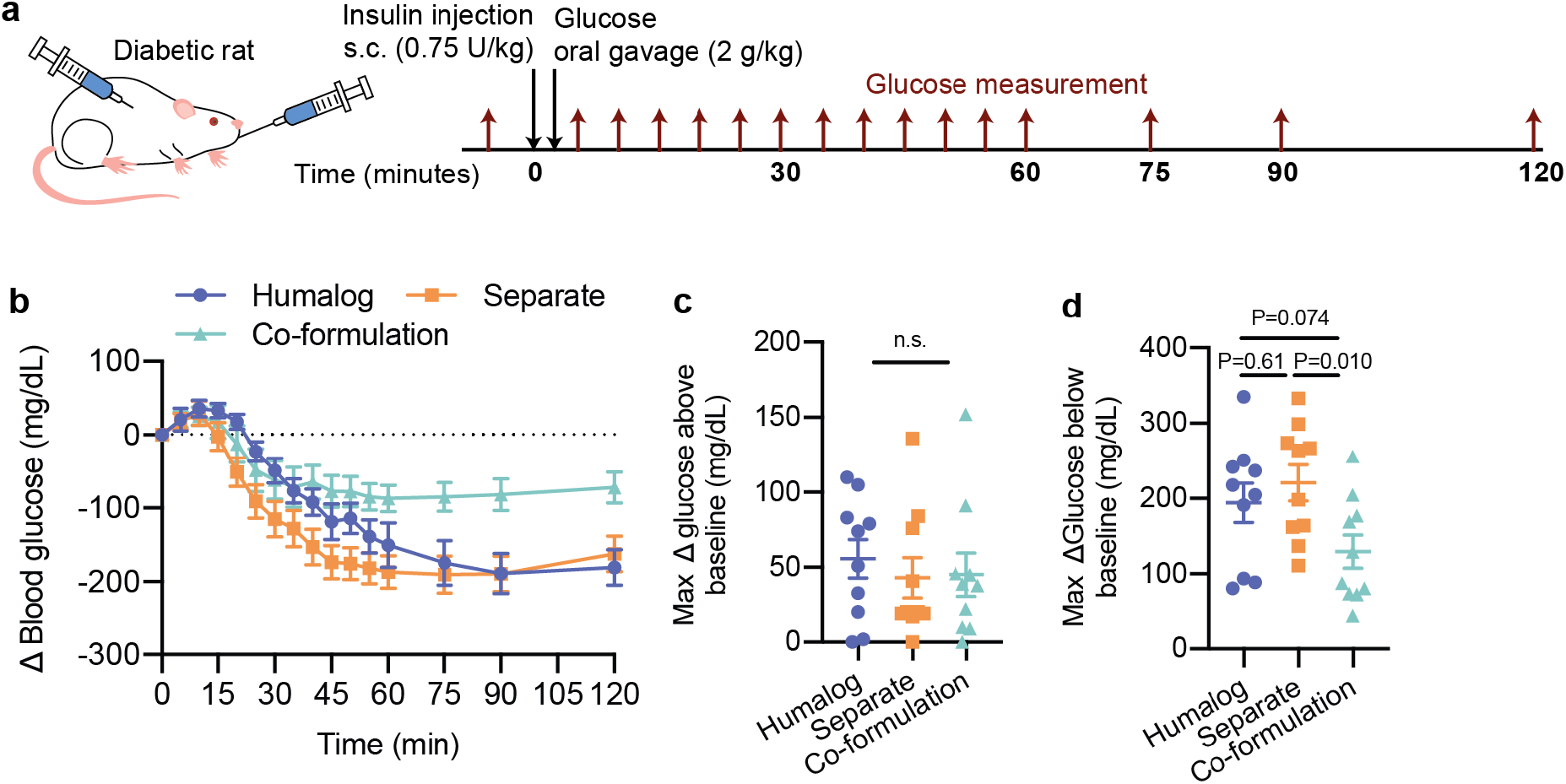
Mealtime simulations with glucose. Fasted male diabetic rats received subcutaneous administration of (i) Humalog, (ii) separate injections of Humalog and pramlintide, or (iii) insulin-pramlintide co-formulation. **a,** Oral glucose challenge where insulin administration (0.75 U/kg) was immediately followed with oral gavage with a glucose solution (2 g/kg). Each rat (n=10) received all treatment groups. **b,** Change in blood glucose after administration is shown. **c,** Max change in glucose above baseline **d,** Max change in glucose below the baseline. All data is shown as mean ± SE. Statistical significance was determined by restricted maximum likelihood repeated measures mixed model. Tukey HSD post-hoc tests were applied to account for multiple comparisons.

## 3. Discussion

In this study, we show that co-formulation of monomeric insulin lispro and pramlintide have ultrafast kinetics with a high degree of overlap resulting in improved glucose management after a glucose challenge. This formulation uses amphiphilic acrylamide copolymer excipient MoNi_23%_ as a stabilizing agent and is physically stable twice as long as commercial Humalog in a stressed aging assay. The pramlintide in the co-formulation results in delayed gastric emptying similar to separately administered pramlintide.

Further, the combined effects of ultrafast insulin and pramlintide delivery synchronized in our co-formulation results in reduced levels of glucose below baseline measurements, while maintaining control of the initial glucose spike in our simulated “mealtime” glucose challenge. The reduced magnitude of glucose levels following administration of the co-formulation is an unexpected, but advantageous, effect since it combines coverage of mealtime glucose spikes with a reduced risk of insulin stacking or post-prandial hypoglycemia. A complete understanding of the complex physiological mechanisms and potential metabolic synergy responsible for this pharmacodynamic effect is challenging or impossible to fully characterize in rats and conducting a large animal study is out of the scope of the present study. While oral glucose tolerance tests are not completely representative of a mixed-meal, slowed gastric emptying is still possible to detect and thus we expect pramlintide to contribute a measurable effect.[16] Based on the results presented in this study, we hypothesize that the shorter duration of insulin action, and resulting greater insulin-pramlintide overlap in the co-formulation, leads to synergy that allows for smoother glucose control. This outcome would suggest that our co-formulation has potential to improve glucose management by reducing the risk of post-prandial hypoglycemia, while reducing patient burden. Further, we hypothesize that greater synergy as a result of synchronized pharmacokinetics could allow for improved glucagon suppression at lower doses as seen in our previous co-formulation study.[8] Future work before translation of this formulation may include better characterization of this effect in other species.

Our data in rats show only trends for increased time to onset (50% of peak up) and time to peak were observed for lispro in the co-formulation compared to Humalog and separate injections. Though, AUC ratios representing the fraction of exposure at various timepoints showed that the co-formulation had a greater fraction of early lispro exposure than separate injections and Humalog up until 30 minutes after injection. These observations are especially exciting because this study was performed in diabetic rats who have much faster insulin absorption rates on account of their loose skin that results in a larger surface area for subcutaneous absorption compared to humans (Figure S5). Indeed, studies comparing rapid-acting insulin analogues and recombinant human insulin, which have distinct differences in time to onset, do not observe differences when compared in rats.[17] Previous study of monomeric lispro in diabetic pigs has shown that time to onset and time to peak are twice as fast for monomeric lispro compared to Humalog.[10] Further, comparison of Humalog, monomeric lispro, and pramlintide kinetics between rats and pigs corroborate previous modeling to suggest the ultrafast kinetics observed here will be conserved across species from rats to humans (Figure S6, S7). Where Humalog time to peak almost doubles from rats (13 ± 1 minutes) to pigs (25 ± 4 minutes), time to peak for monomeric lispro (delivered as part of the co-formulation in rats) is similar in both species (11 ± 1 minutes in rats and 9 ± 2 minutes in pigs) (Figure S7).[10] The conservation of time to peak exposure from rats to pigs is highly promising for the translation of these ultrafast insulin kinetics to human trials and would result in kinetics faster than current commercial formulations (Figure S7).

Beyond improved bolus insulin delivery using the co-formulation, delivering an insulin with these ultrafast kinetics synchronously with pramlintide presents opportunities for applications in insulin infusion pumps and “artificial pancreas” closed-loop systems. Studies using two separate pumps delivering insulin and pramlintide at a fixed ratio have shown that dual-hormone replacement results in reduced mean glucose compared to insulin alone.[18] Recently, this two-pump delivery approach has been used in a closed-loop system and an increased time in target glucose range was observed for patients who received a fixed ratio of rapid-acting insulin and pramlintide compared to rapid-acting insulin alone.[19] A stable insulin-pramlintide co-formulation would enable the implementation of this dual-hormone treatment in closed-loop systems outside of clinical trials where using two separate infusion pumps is impractical. The synchronized insulin-pramlintide kinetics and shorter duration of insulin action in our co-formulation also have future promise for better autonomous insulin delivery. At present, these closed-loop systems require patients to input carbohydrates counts at mealtimes and are not fully autonomous, in part because insulin absorption kinetics are not rapid enough to reduce mealtime glucose excursions, and the extended duration of insulin action can result in “insulin stacking” leading to post-prandial hypoglycemia. An ultrafast insulin-pramlintide co-formulation has the potential to rapidly react to mealtime spikes, as the insulin will have immediate onset and the pramlintide will slow the appearance of glucose (through delayed gastric emptying). Further, with shorter duration of insulin action, the risk of hypoglycemia, as a result of insulin stacking would be reduced.

As MoNi_23%_ is a new excipient, future work will have to complete robust safety and biocompatibility tests before translation to humans. Preliminary cytotoxicity and biocompatibility studies suggest MoNi_23%_ is well tolerated, and adverse effects are not anticipated with its use.[10] An additional area of investigation for future studies is the chemical stability of our co-formulation. We have demonstrated pramlintide in our co-formulation is physically stable under stressed-aging conditions for longer durations that current commercial Humalog, and that it is active *in vivo*, demonstrating delayed gastric emptying after administration. Though, before commercialization, the chemical stability of our formulation will have to be investigated to ensure formulation integrity over a long shelf-life.

## 4. Conclusion

Together, these studies demonstrate that a stable insulin-pramlintide co-formulation drug product candidate utilizing monomeric insulin exhibits synchronized ultrafast insulin-pramlintide pharmacokinetics that result in better glycemic control in a mealtime simulation. This co-formulation has potential to improve glucose management and reduce patient burden in clinical applications using it for both direct bolus administration as well as in insulin infusion pumps or artificial pancreas closed-loop systems. While we focus on the treatment of type I diabetes in this study, anyone taking insulin therapies, including patients with type II diabetes, would benefit from a single administration, dual-hormone drug product such as this.

## 5. Methods

### 5.1 Materials

Our amphiphilic acrylamide copolymer excipient acryloylmorpholine_77%_-N-isopropylacrylamide_23%_(MoNi_23%_) was prepared according to published protocols.[10] Characterization of MoNi_23%_ molecular weight and monomer composition can be found in Table S1. in Humalog (Eli Lilly) and pramlintide (BioTang) were purchased and used as received. For zinc-free lispro, Zinc(II) was removed from the insulin lispro through competitive binding by addition of ethylenediaminetetraacetic acid (EDTA), which exhibits a dissociation binding constant approaching attomolar concentrations (K_D_~10^−18^ M).[20] EDTA was added to formulations (4 eq with respect to zinc) to sequester zinc from the formulation and then lispro was isolated using PD MidiTrap G-10 gravity columns (GE Healthcare) to buffer exchange into water. The solution was then concentrated using Amino Ultra 3K centrifugal units (Millipore) and reformulated with 2.6 wt.% glycerol, 0.85 wt.% phenoxyethanol in 10 mM phosphate buffer (pH=7.4). All other reagents were purchased from Sigma-Aldrich unless otherwise specified.

### 5.2 Methods

#### SEC-MALS

Insulin association state composition for monomeric insulin formulation was obtained using SEC-MALS as previously reported.[13] Zinc-free insulin lispro was evaluated in a buffer containing glycerol (2.6%) and phenoxyethanol (0.85%). Briefly, number-averaged molecular weight (MW) and dispersity (Đ = Mw/Mn) of formulations were obtained using size exclusion chromatography (SEC) carried out using a Dionex Ultimate 3000 instrument (including pump, autosampler, and column compartment) outfitted with a Dawn Heleos II Multi Angle Light Scattering detector, and a Optilab rEX refractive index detector. The column was a Superose 6 Increase 10/300 GL from GE healthcare. Data was analyzed using Astra 6.0 software. The fraction of each insulin association state was derived by fitting the experimentally derived number-average and weight-average molecular weights to Equation 1 and Equation 2 below. *m, d* and *h*, respectively, represent the molar fractions of monomeric, dimeric and hexameric insulin while *I* represents the molecular weight of monomeric insulin lispro. The solver was constrained so that *m*+*d*+*h*=1 while *m*, *d* and *h* remain between 0 and 1.

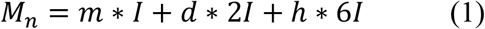

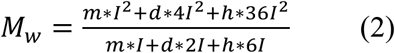

#### In vitro stability

Aggregation assays used to evaluate stability were adapted from Webber et al.[15] Briefly, formulations were aliquoted 150 μL per well (n = 3/group) in a clear 96-well plate and sealed with optically clear and thermally stable seal (VWR). The plate was incubated in a microplate reader (BioTek SynergyH1 microplate reader) at 37 °C with continuous agitation (567 cpm). Absorbance readings were taken every 10 minutes at 540 nm for the duration of the experiment. The formation of insulin or pramlintide aggregates leads to light scattering and a reduction in the transmittance of samples (time to aggregation = time to 10% change in transmittance). Controls included: (i) Humalog (100 U/mL), (ii) Humalog (100U/mL) + Pramlintide (1:6 lispro:pramlintide), (iii) zinc-free lispro (100U/mL lispro, 2.6 wt.% glycerol, 0.85 wt.% phenoxyethanol, pH=7.4). The stability of an insulin-pramlintide co-formulation (100U/mL lispro, 1:6 lispro:pramlintide, 2.6 wt.% glycerol, 0.85 wt.% phenoxyethanol, pH=7.4) mixed with 0.1 mg/mL MoNi_23%_was evaluated.

#### Streptozotocin induced model of diabetes in rats

Male Sprague Dawley rats (Charles River) were used for experiments. Animal studies were performed in accordance with the guidelines for the care and use of laboratory animals; all protocols were approved by the Stanford Institutional Animal Care and Use Committee (Protocol #32873). The protocol used for streptozotocin (STZ) induction adapted from the protocol by Kenneth K. Wu and Youming Huan and has been previously reported.[8, 12–13, 21] Briefly, male Sprague Dawley rats 160-230g (8-10 weeks) were weighed and fasted in the morning 6-8 hours prior to treatment with STZ. STZ was diluted to 10mg/mL in the sodium citrate buffer immediately before injection. STZ solution was injected intraperitoneally at 65mg/kg into each rat. Rats were provided with water containing 10% sucrose for 24 hours after injection with STZ. Rat blood glucose levels were tested for hyperglycemia daily after the STZ treatment via tail vein blood collection using a handheld Bayer Contour Next glucose monitor (Bayer). Diabetes was defined as having 3 consecutive blood glucose measurements >400 mg/dL in non-fasted rats.

#### In vivo pharmacokinetics and pharmacodynamics in diabetic rats

Diabetic rats were fasted for 4-6 hours before injection. For pharmacokinetic experiments rats were injected with 1U insulin formulation (~2 U/kg) followed immediately (< 30 seconds after injection) by oral gavage with 1g/kg glucose solution. The dose of insulin was chosen because it could be tolerated by the rats, and allowed for delivery with an insulin syringe with minimal dilution. Formulations tested were: (i) Humalog, (ii) separate injections of Humalog and pramlintide (1:6 pramlintide:lispro, pH=4), (iii) insulin-pramlintide co-formulation (100 U/mL lispro, 1:6 lispro:pramlintide, 2.6 wt.% glycerol, 0.85 wt.% phenoxyethanol, 0.1 mg/mL MoNi_23%_, pH=7.4). A cohort of 11 rats each received each formulation once, and the order the formulations were given in was randomized. To allow for accurate dosing and to avoid dilution effects (dilution favours the insulin monomer) formulations were diluted two-fold (10 μL formulation + 10 μL formulation buffer) immediately before administration. After injection, blood glucose measurements were taken using a handheld glucose monitor (Bayer Contour Next) and additional blood was collected (Sarstedt serum tubes) for analysis with ELISA. Timepoints were taken every 3 minutes for the first 30 minutes, then every 5 minutes for the next 30 minutes, then at 75, 90, and 120 minutes. Serum pramlintide concentrations were quantified using a human amylin ELISA kit (Millipore Sigma). Serum lispro concentrations were quantified using Northern Lights Mercodia Lispro NL-ELISA. A second pharmacodynamics experiment was performed to try to better match insulin dose with oral glucose dose to better simulate meal-time glucose management. The same formulations were tested but doses were changed to 0.75 U/kg insulin delivered subcutaneously immediately before oral gavage with 2 g/kg glucose. The lower dose was chosen to try to better match the carbohydrate load, however this was limited by the volume of undiluted insulin that could be practically administered to the rats and thus the insulin dose still resulted in a net decrease in glucose levels. A 10 μL Hamilton syringe was used to allow accurate dosing of undiluted (100 U/mL) formulations. A cohort of 10 rats each received each formulation once, and the order the formulations were given in was randomized. Only glucose was measured and timepoints were taken every 5 minutes for the first hour, followed by measurements at 75, 90 and 120 minutes.

#### Gastric emptying in diabetic rats

Acetaminophen is used as a model compound to evaluate gastric emptying at mealtimes. Diabetic rats were fasted for 4-6 hours before experiment start. Rats were then injected subcutaneously with one of the following formulations (2 U/kg): (i) Humalog, (ii) separate injections of Humalog and pramlintide (1:6 pramlintide:lispro, pH=4), (iii) insulin-pramlintide co-formulation (100 U/mL lispro, 1:6 lispro:pramlintide, 2.6 wt.% glycerol, 0.85 wt.% phenoxyethanol, 0.1 mg/mL MoNi_23%_, pH=7.4). To allow for accurate dosing and to avoid dilution effects (dilution favours the insulin monomer) formulations were diluted two-fold (10 μL formulation + 10 μL formulation buffer) immediately before administration. A cohort of 11 rats each received each formulation once, and the order the formulations were given in was randomized. Acetaminophen was administered via oral gavage as a slurry in phosphate buffer (100 mg/kg) immediately after insulin administration. (Tips of feeding tubes were dipped in glucose solution before oral gavage to reduce stress of administration).[22] Blood samples were collected for ELISA (Neogen) at −30, 0, 15, 30, 60, 90, 120, ands 150 minutes after injection.

#### Statistics

All results are expressed as mean ± standard error (SE) unless specified otherwise. Sample size for each experiment is included in the corresponding methods section as well as figure captions. All statistical analyses were performed as general linear models in JMP Pro version 14. Comparisons between formulations were conducted using the restricted maximum likelihood repeated measures mixed model. Post-hoc Tukey HSD tests for multiple comparisons was applied when formulation was a significant fixed effect, and adjusted p-values were reported. Rat was included as a variable in the model as a random effect blocking (control) factor to account for individual variation in rat responses. (Each rat received every formulation and acted as its own control). Statistical significance was considered as P < 0.05. For Fig. 2h-l, post-hoc Bonferroni correction was applied to account for comparison of formulations at multiple exposure timepoints (In addition to Tukey HSD correction) and significance was adjusted to α=0.01.

## Supporting Information

Supporting Information is available from the Wiley Online Library or from the author.

## Acknowledgments

This work was funded in part by NIDDK R01 (NIH grant #R01DK119254), and a Pilot and Feasibility funding from the Stanford Diabetes Research Center (NIH grant #P30DK116074), as well as the American Diabetes Association Grant (1-18-JDF-011) and a Research Starter Grant from the PhRMA. Support is also provided by the Stanford Maternal and Child Health Research Institute through the SPARK Translational Research Program. C.L.M. was supported by the NSERC Postgraduate Scholarship and the Stanford BioX Bowes Graduate Student Fellowship. A.I.D. was supported by the Schmidt Science Fellows Award. J.L.M was supported Department of Defense NDSEG Fellowship and by a Stanford Graduate Fellowship. The authors thank the Stanford Animal Diagnostic Lab and the Veterinary Service Centre staff for their technical assistance.

## Author Contributions

C.L.M, P.C.C. and E.A.A designed the experiments. C.L.M., A.I.D., E.T.V., L.T.N. performed the experiments. J.L.M. synthesized the polymers. C.L.M., P.C.C., J.L.M analyzed the data. C.L.M. and E.A.A. wrote the manuscript. All authors revised the manuscript.

## Data Availability

All data supporting the results in this study are available within the Article and its Supplementary Information. Raw data files are available from the corresponding author upon reasonable request.

## References

[1] D. L. Hay, S. Chen, T. A. Lutz, D. G. Parkes, J. D. Roth, Pharmacol. Rev. 2015, 67 (3), 564, https://doi.org/10.1124/pr.115.010629.

[2] C. Martin, Diabetes Educ. 2006, 32 (Supplement 3), 101S, https://doi.org/10.1177/0145721706288237.

a) R. Ratner, M. S. Whitehouse F Fau - Fineman, S. Fineman Ms Fau - Strobel, L. Strobel S Fau - Shen, D. G. Shen L Fau - Maggs, O. G. Maggs Dg Fau - Kolterman, C. Kolterman Og Fau - Weyer, C. Weyer, Exp. Clin. Endocrinol. Diabetes 2005, 113 (0947-7349 (Print)), 199;

b) R. E. Ratner, R. Dickey, M. Fineman, D. G. Maggs, L. Shen, S. A. Strobel, C. Weyer, O. G. Kolterman, Diabet. Med. 2004, 21 (11), 1204;

c) F. Whitehouse, D. F. Kruger, M. Fineman, L. Shen, J. A. Ruggles, D. G. Maggs, C. Weyer, O. G. Kolterman, Diabetes Care 2002, 25 (4), 724;

d) G. J. Ryan, L. J. Jobe, R. Martin, Clin.Ther. 2005, 27 (10), 1500, https://doi.org/10.1016/j.clinthera.2005.10.009;

e) S. Edelman, S. Garg, J. Frias, D. Maggs, Y. Wang, B. Zhang, S. Strobel, K. Lutz, O. Kolterman, Diabetes Care 2006, 29 (10), 2189, https://doi.org/10.2337/dc06-0042;

f) L. M. Rodriguez, K. J. Mason, M. W. Haymond, R. A. Heptulla, Pediatr. Res. 2007, 62 (6), 746, https://doi.org/10.1203/PDR.0b013e318159af8c;

g) S. A. Weinzimer, J. L. Sherr, E. Cengiz, G. Kim, J. L. Ruiz, L. Carria, G. Voskanyan, A. Roy, W. V. Tamborlane, Diabetes Care 2012, 35 (10), 1994, https://doi.org/10.2337/dc12-0330.

[4] C. Hampp, D. G. Borders-Hemphill V Fau - Moeny, D. K. Moeny Dg Fau - Wysowski, D. K. Wysowski, Use of antidiabetic drugs in the U.S., 2003-2012. Diabetes Care. 2014, 37, 1367–74.

[5] H. Wang, A. Abedini, B. Ruzsicska, D. P. Raleigh, Biochemistry 2014, 53 (37), 5876, https://doi.org/10.1021/bi500592p.

[6] J. Koda, M. Fineman, T. Rink, G. Dailey, D. Muchmore, L. Linarelli, Lancet 1992, 339 (8802), 1179, https://doi.org/10.1016/0140-6736(92)90785-2.

a) C. Mathieu, P. Gillard, K. Benhalima, Nat. Rev. Endocrinol. 2017, 13 (7), 385, https://doi.org/10.1038/nrendo.2017.39;

b) F. Holleman, J. B. L. Hoekstra, N. Engl. J. Med. 1997, 337 (3), 176, https://doi.org/10.1056/NEJM199707173370307;

c) K. Gast, A. Schüler, M. Wolff, A. Thalhammer, H. Berchtold, N. Nagel, G. Lenherr, G. Hauck, R. Seckler, Pharm. Res. 2017, 34 (11), 2270, https://doi.org/10.1007/s11095-017-2233-0.

[8] C. L. Maikawa, A. A. A. Smith, L. Zou, G. A. Roth, E. C. Gale, L. M. Stapleton, S. W. Baker, J. L. Mann, A. C. Yu, S. Correa, A. K. Grosskopf, C. S. Liong, C. M. Meis, D. Chan, M. Troxell, D. M. Maahs, B. A. Buckingham, M. J. Webber, E. A. Appel, Nat. Biomed. Eng. 2020, 4 (5), 507, https://doi.org/10.1038/s41551-020-0555-4.

a) M. C. Riddle, K. C. J. Yuen, T. W. de Bruin, K. Herrmann, J. Xu, P. Öhman, O. G. Kolterman, Diabetes Obes. Metab. 2015, 17 (9), 904, https://doi.org/10.1111/dom.12504;

b) O. G. Kolterman, S. Schwartz, C. Corder, B. Levy, L. Klaff, J. Peterson, A. Gottlieb, Diabetologia 1996, 39 (4), 492, https://doi.org/10.1007/BF00400683;

c) J. Plank, A. Wutte, G. Brunner, A. Siebenhofer, B. Semlitsch, R. Sommer, S. Hirschberger, T. R. Pieber, Diabetes Care 2002, 25 (11), 2053, https://doi.org/10.2337/diacare.25.11.2053;

d) R. J. Pettis, L. Hirsch, C. Kapitza, L. Nosek, U. Hövelmann, H.-J. Kurth, D. E. Sutter, N. G. Harvey, L. Heinemann, Diabetes Technol. Ther. 2011, 13 (4), 443, https://doi.org/10.1089/dia.2010.0183;

e) G. Andersen, G. Meiffren, D. Lamers, J. H. DeVries, A. Ranson, C. Seroussi, B. Alluis, M. Gaudier, O. Soula, T. Heise, Diabetes Obes. Metab. 2018, https://doi.org/10.1111/dom.13442;

f) H. Linnebjerg, Q. Zhang, E. LaBell, M. A. Dellva, D. E. Coutant, U. Hövelmann, L. Plum-Mörschel, T. Herbrand, J. Leohr, Clin. Pharmacokinet. 2020, 59 (12), 1589, https://doi.org/10.1007/s40262-020-00903-0.

[10] J. L. Mann, C. L. Maikawa, A. A. A. Smith, A. K. Grosskopf, S. W. Baker, G. A. Roth, C. M. Meis, E. C. Gale, C. S. Liong, S. Correa, D. Chan, L. M. Stapleton, A. C. Yu, B. Muir, S. Howard, A. Postma, E. A. Appel, Sci. Transl. Med. 2020, 12, eaba6676.

a) V. Sluzky, A. M. Klibanov, R. Langer, Biotechnol. Bioeng. 1992, 40 (8), 895, https://doi.org/10.1002/bit.260400805;

b) V. Sluzky, J. A. Tamada, A. M. Klibanov, R. Langer, Proc. Natl. Acad. Sci. U. S. A. 1991, 88 (21), 9377, https://doi.org/10.1073/pnas.88.21.9377;

c) L. Nault, P. Guo, B. Jain, Y. Bréchet, F. Bruckert, M. Weidenhaupt, Acta Biomater. 2013, 9 (2), 5070, https://doi.org/10.1016/j.actbio.2012.09.025.

[12] C. L. Maikawa, J. L. Mann, A. Kannan, C. M. Meis, A. K. Grosskopf, B. S. Ou, A. A. A. Autzen, G. G. Fuller, D. M. Maahs, E. A. Appel, Biomacromolecules 2021, https://doi.org/10.1021/acs.biomac.1c00474.

[13] C. L. Maikawa, A. A. A. Smith, L. Zou, C. M. Meis, J. L. Mann, M. J. Webber, E. A. Appel, Adv. Ther. 2019, 75, 1900094, https://doi.org/10.1002/adtp.201900094.

a) T. Sanke, T. Hanabusa, Y. Nakano, C. Oki, K. Okai, S. Nishimura, M. Kondo, K. Nanjo, Diabetologia 1991, 34 (2), 129;

b) F. Micheletto, C. Dalla Man, O. Kolterman, E. Chiquette, K. Herrmann, J. Schirra, B. Kovatchev, C. Cobelli, Diabetes Technol. Ther. 2013, 15 (10), 802, https://doi.org/10.1089/dia.2013.0054.

[15] M. J. Webber, E. A. Appel, B. Vinciguerra, A. B. Cortinas, L. S. Thapa, S. Jhunjhunwala, L. Isaacs, R. Langer, D. G. Anderson, Proc. Natl. Acad. Sci. U. S. A. 2016, 113 (50), 14189, https://doi.org/10.1073/pnas.1616639113.

a) A. A. Young, W. Vine, B. R. Gedulin, R. Pittner, S. Janes, L. S. L. Gaeta, A. Percy, C. X. Moore, J. E. Koda, T. J. Rink, K. Beaumont, Drug Dev. Res. 1996, 37 (4), 231, https://doi.org/10.1002/(SICI)1098-2299(199604)37:4

b) M. Sala-Rabanal, C. Ghezzi, B. A. Hirayama, V. Kepe, J. Liu, J. R. Barrio, E. M. Wright, J. Physiol. 2018, 596 (13), 2473, https://doi.org/10.1113/JP275934.

[17] A. Plum, H. Agersø, L. Andersen, Drug Metab.Dispos. 2000, 28 (2), 155.

[18] M. C. Riddle, R. Nahra, J. Han, J. Castle, K. Hanavan, M. Hompesch, D. Huffman, P. Strange, P. Öhman, Diabetes Care 2018, https://doi.org/10.2337/dc18-1091.

[19] A. Haidar, M. A. Tsoukas, S. Bernier-Twardy, J.-F. Yale, J. Rutkowski, A. Bossy, E. Pytka, A. El Fathi, N. Strauss, L. Legault, Diabetes Care 2020, 43 (3), 597, https://doi.org/10.2337/dc19-1922.

a) G. Berthon, Handbook Of Metal-ligand Interactions in Biological Fluids: Bioinorganic chemistry, Marcel dekker, New York 1995;

b) R. S. Waters, N. A. Bryden, K. Y. Patterson, C. Veillon, R. A. Anderson, Biol. Trace Elem. Res. 2001, 83 (3), 207, https://doi.org/10.1385/bter:83:3:207.

[21] K. K. Wu, Y. Huan, Curr. Protoc. Pharmacol. 2008, (1934-8290 (Electronic)), 5.47.1.

[22] A. F. Hoggatt, J. Hoggatt, M. Honerlaw, L. M. Pelus, J. Am. Assoc. Lab Anim. Sci. 2010, 49 (3), 329.

[23] C. L. Maikawa, A. A. A. Smith, L. Zou, G. A. Roth, E. C. Gale, L. M. Stapleton, S. W. Baker, J. L. Mann, A. C. Yu, S. Correa, A. K. Grosskopf, C. S. Liong, C. M. Meis, D. Chan, M. D. Troxell, D. M. Maahs, B. A. Buckingham, M. J. Webber, E. A. Appel, Nat. Biomed. Eng. 2020, 4, 507.

a) L. Heinemann, T. Heise, L. C. Wahl, M. E. Trautmann, J. Ampudia, A. A. R. Starke, M. Berger, Diabet. Med. 1996, 13 (7), 625, https://doi.org/10.1002/(SICI)1096-9136(199607)13:7

b) A. Lindholm, J. McEwen, A. P. Riis, Diabetes Care 1999, 22 (5), 801, https://doi.org/10.2337/diacare.22.5.801;

c) K. Rave, O. Klein, A. D. Frick, R. H. A. Becker, Diabetes Care 2006, 29 (8), 1812, https://doi.org/10.2337/dc06-0383.

a) M. Fath, T. Danne, T. Biester, L. Erichsen, O. Kordonouri, H. Haahr, Pediatr. Diabetes 2017, 18 (8), 903, https://doi.org/10.1111/pedi.12506;

b) T. Heise, T. R. Pieber, T. Danne, L. Erichsen, H. Haahr, Clin. Pharmacokinet. 2017, 56 (5), 551, https://doi.org/10.1007/s40262-017-0514-8.

[26] M. Shiramoto, R. Nasu, T. Oura, M. Imori, K. Ohwaki, J. Diabetes Investig. 2020, 11 (3), 672, https://doi.org/10.1111/jdi.13195.

