## Supplemental Information for "Ultra-fast insulin-pramlintide co-formulation for improved glucose management in diabetic rats"

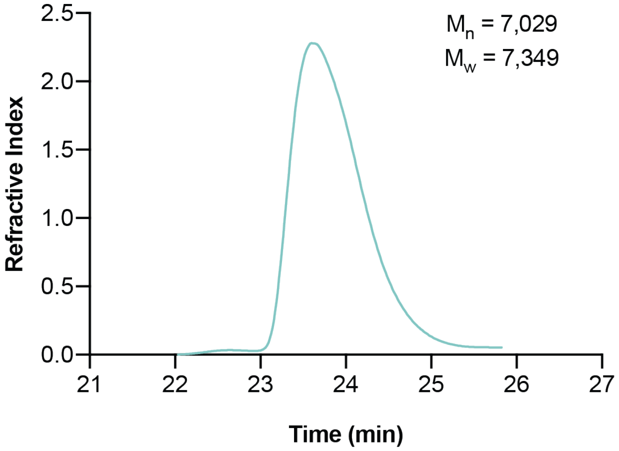


**Figure S1. SEC-MALS elution curve.** SEC-MALS elution profiles of the distribution of insulin lispro aggregation states with number-averaged molecular weight and weight-averaged molecular weight. Used to calculate percentage of insulin association states. Number-averaged molecular weight (MW) and dispersity (Đ = Mw/Mn) of formulations were obtained using size exclusion chromatography (SEC) carried out using a Dionex Ultimate 3000 instrument (including pump, autosampler, and column compartment) outfitted with a Dawn Heleos II Multi Angle Light Scattering detector, and a Optilab rEX refractive index detector. The column was a Superose 6 Increase 10/300 GL from GE healthcare. Data was analyzed using Astra 6.0 software.

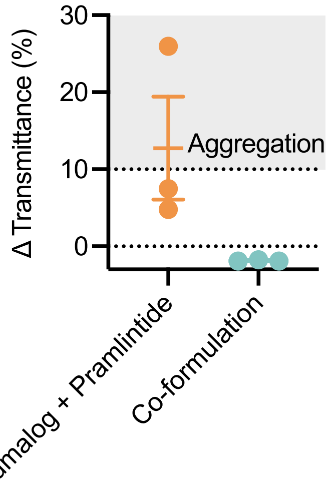


**Figure S2. Initial transmittance for Humalog + pramlintide and co-formulation.** Initial transmittance values for (i) Humalog + pramlintide control group and (ii) co-formulation in the stability assay (Figure 1f) shown as the change in transmittance of (Average initial Humalog transmittance) - formulation initial transmittance. The decreased transmittance observed for the the Humalog + Pramlintide samples before the aging study indicates that there is poor solubility when these two formulations are mixed. In comparison, the initial transmittance for the co-formulation is optically clear to the eye and shows little difference from initial Humalog transmittance.

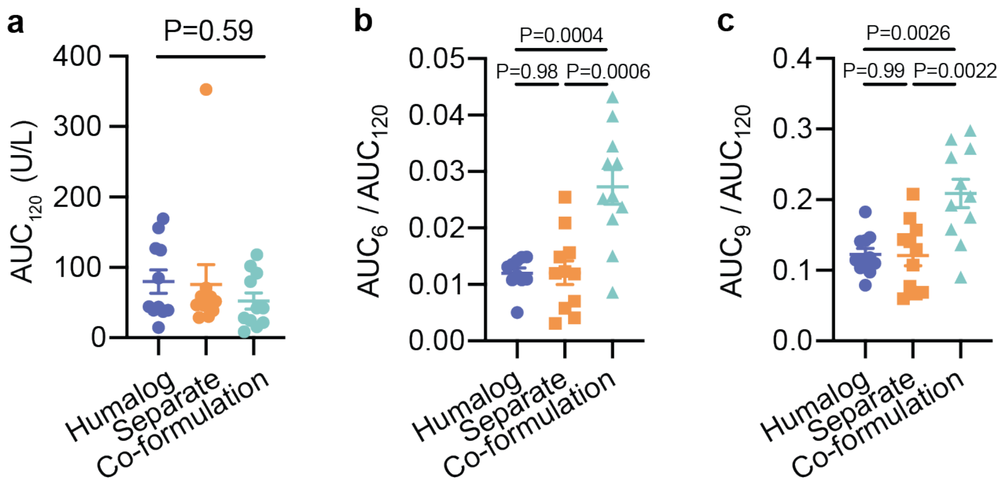


**Figure S3. Lispro area under the curve and exposure ratios. a,** Total lispro exposure as area under the pharmacokinetic curve. **b, c,** Fraction of lispro exposure as a ratio of $AUC_t$/ $AUC_{120}$ at **b,** t=3; **c,** t=9. Statistical significance was determined by restricted maximum likelihood repeated measures mixed model. Tukey HSD post-hoc tests were applied to account for multiple comparisons (b,c).

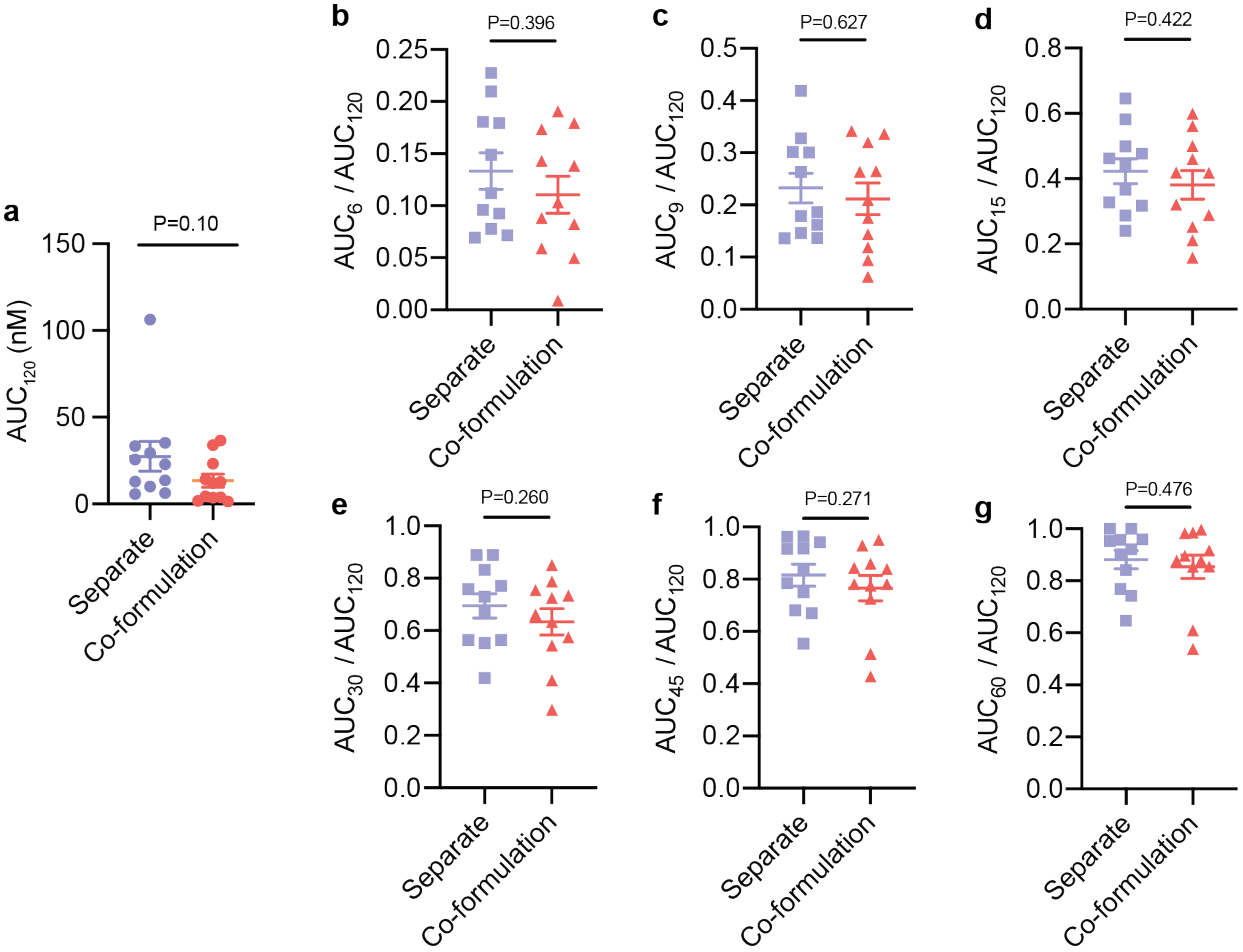


**Figure S4. Pramlintide area under the curve and exposure ratios. a,** Total pramlintide exposure as area under the pharmacokinetic curve. **b-f,** Fraction of pramlintide exposure as a ratio of $AUC_t/AUC_{120}$ at **b,** t=6; **c,** t=9; **d,** t=15; **e,** t=30; **f,** t=45; **g,** t=60. Statistical significance was determined by restricted maximum likelihood repeated measures mixed model.

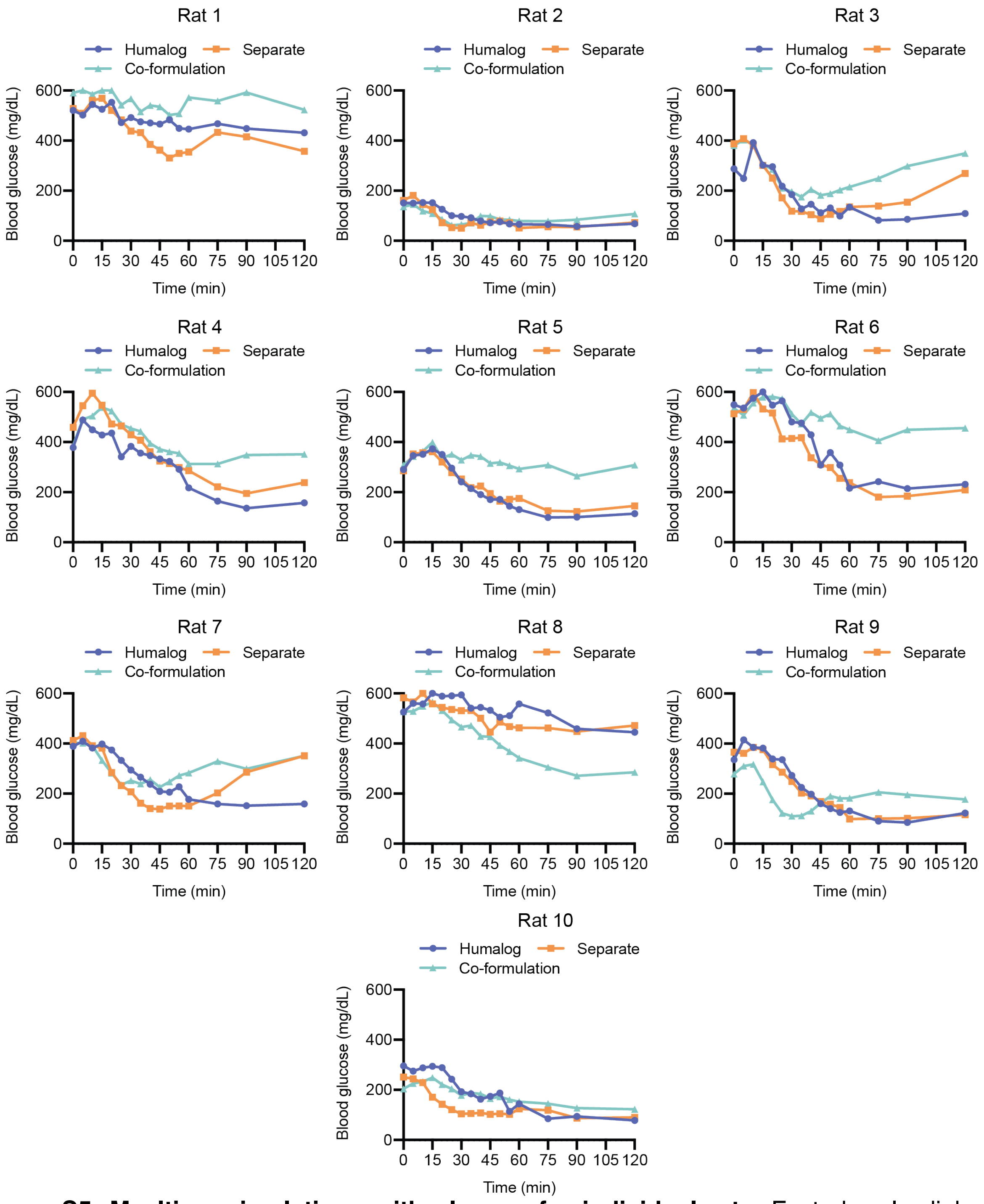


**Figure S5. Mealtime simulations with glucose for individual rats.** Fasted male diabetic rats received subcutaneous administration of (i) Humalog, (ii) separate injections of Humalog and pramlintide, or (iii) insulin-pramlintide co-formulation. Blood glucose measurements are shown for each rat after oral glucose challenge where insulin administration (0.75 U/kg) was immediately followed with oral gavage with a glucose solution (2 g/kg). Each rat (n=10) received all treatment groups and the order that treatments were given was randomized. When rats were fasted the morning of the experiment (5-6 hours before insulin administration), rats were given insulin to adjust glucose levels to be within a measurable range (<600 mg/dL) and ideally 300-400 mg/dL. Differing basal glucose secretion levels and the degree to which each rat was diabetic led to variable success of adjusting glucose levels within this range and thus the goal was adjusted to instead try to have each individual rat start at a similar glucose level for all three formulations.

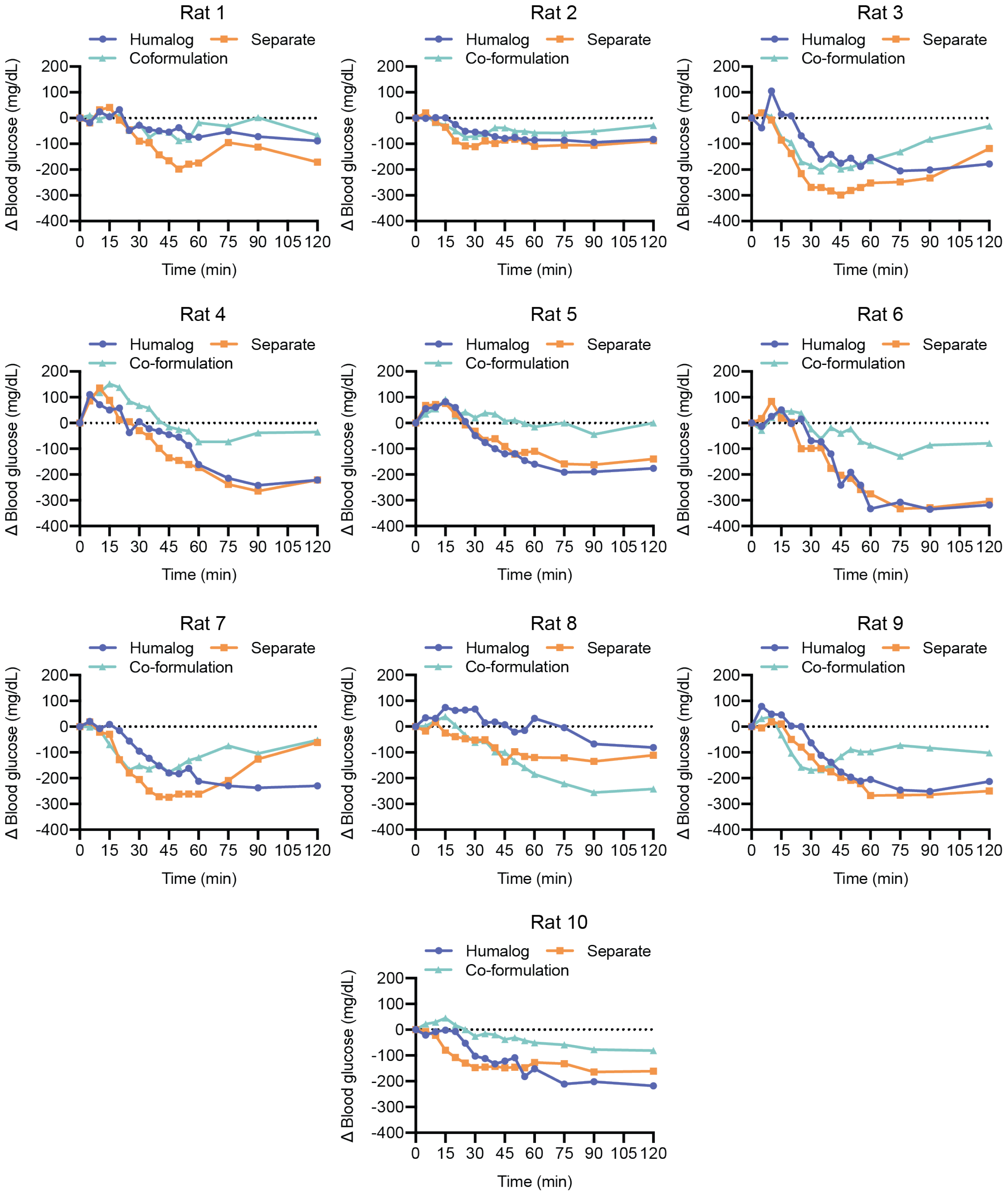


**Figure S6. Mealtime simulations with glucose for individual rats.** Fasted male diabetic rats received subcutaneous administration of (i) Humalog, (ii) separate injections of Humalog and pramlintide, or (iii) insulin-pramlintide co-formulation. Change in glucose measurements for each individual rat are shown after oral glucose challenge where insulin administration (0.75 U/kg) was immediately followed with oral gavage with a glucose solution (2 g/kg) (Baseline corrected data of Figure S5). Each rat (n=10) received all treatment groups and the order that treatments were given was randomized.

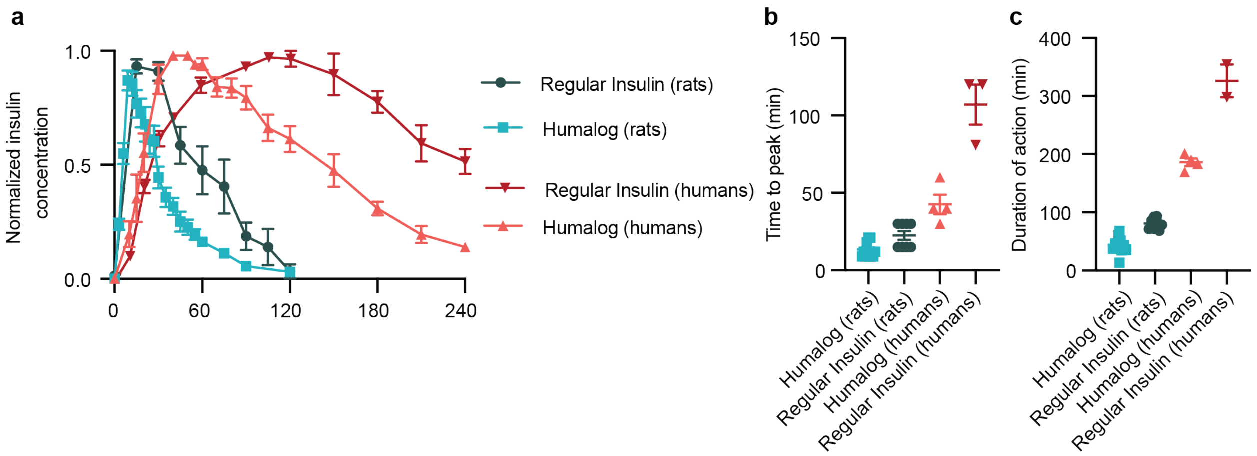


**Figure S7 Interspecies pharmacokinetics for Humalog and Humulin.** a, Normalized pharmacokinetics for commercial Humalog and regular human insulin (ex. Humulin R) delivered in rats and humans. b, Time to peak exposure for each formulation and c, duration of action for each formulation. Differences in time to onset and time to peak in rats between rapid-acting and regular insulin formulations are minimal and difficult to detect. However, in humans, these small differences translate to distinct differences in time to onset and time to peak. This suggests that the trend for more rapid action we observe for insulin in our co-formulation compared to Humalog could translate into substantial differences in humans. Duration of action is defined here as peak width at 25% peak height ($time_{25\%\ down}$ - $time_{25\%\ up}$). Rat data for Humalog is taken from this study, and rat data for regular human insulin is adapted from previous work. Human Humalog data is from three external studies and has been adapted from presentation in previous work.[1-4] Human regular human insulin data is from three external studies.[5-7]

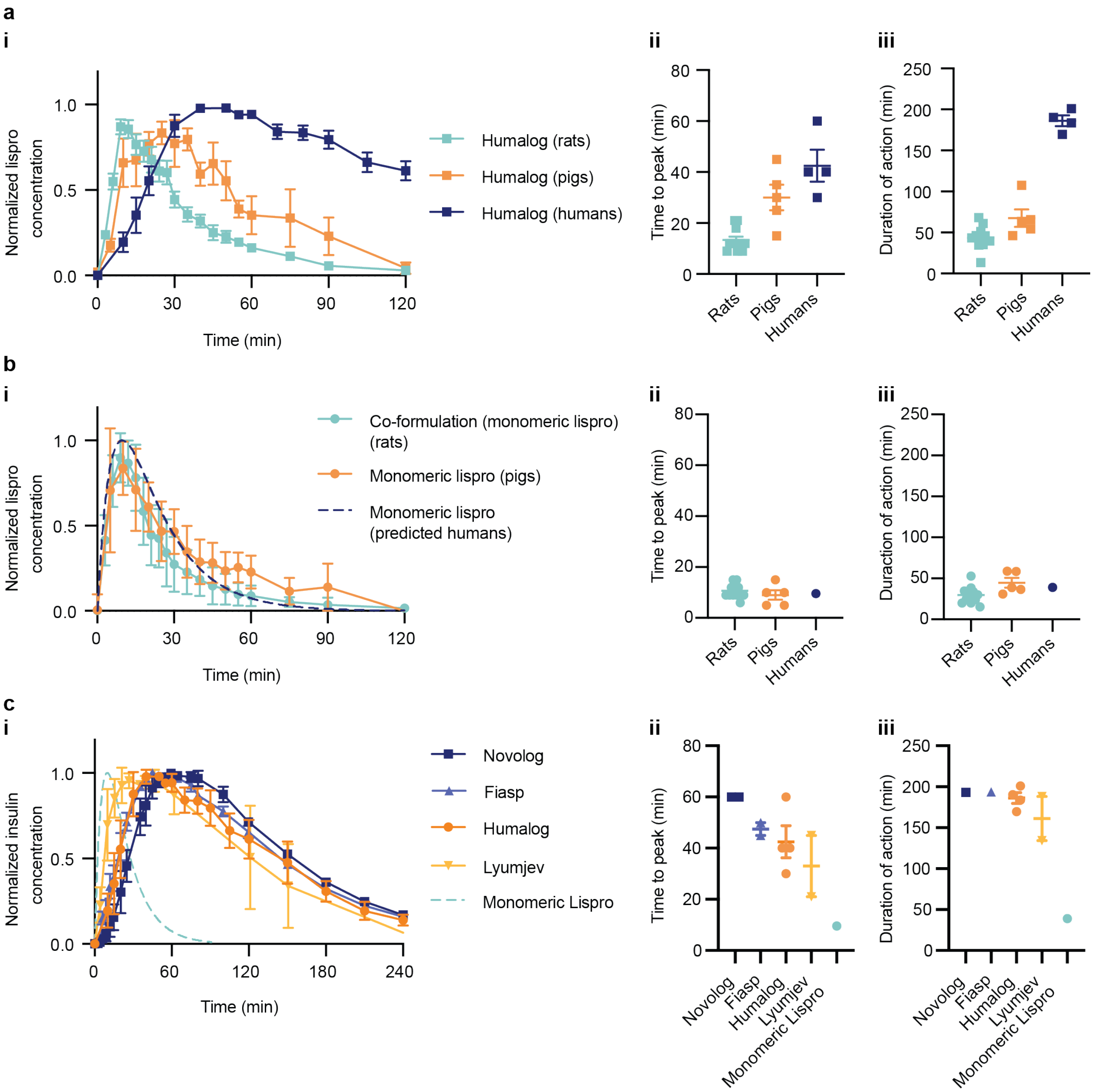


**Figure S8. Interspecies pharmacokinetics for Humalog and monomeric lispro formulations. a,** (i) Normalized pharmacokinetics for commercial Humalog delivered in rats, pigs and humans, (ii) time to peak, and (iii) duration of action. **b,** (i) Normalized pharmacokinetics for monomeric insulin delivered in rats, pigs and humans, (ii) time to peak, and (iii) duration of action. Humalog shows increased time to onset and longer duration of action as you shift to species with more complex subcutaneous architecture (rats < pigs < humans). In contrast, monomeric lispro has very similar onset and duration of action between rats and pigs. Since the difference between rats and pigs is minimal for this formulation, it is also likely that the difference between pigs and humans will be small. Further, we observe that pramlintide - which only exists as a monomer - has very similar kinetics between pigs and humans. **c,** (i) Normalized pharmacokinetics for commercial formulations in humans, (ii) time to peak exposure for each formulation and (iii) duration of action for each formulation. Even with the shift towards faster time to peak with next generation rapid-acting insulins like Fiasp and Lyumjev there have not been similar increases in reducing duration of action. Duration of action is defined here as peak width at 25% peak height ($time_{25\%\ down}$ - $time_{25\%\ up}$). **(a,b)** Rat data is taken from this study (monomeric lispro in rats was delivered as part of the co-formulation). Pig data was adapted from previous work.[8] Human Humalog data is from four external studies.[1-4] Predicted monomeric lispro in humans has been adapted from pharmacokinetic modeling from our previous work in pigs.[8] **(c)** Data is adapted from clinical studies in humans for (i) regular human insulin,[5-7] (ii) Novolog (Novo Nordisk),[9, 10] (iii) Fiasp (Novo Nordisk),[9, 10] (iv) Humalog (Eli Lily),[1-4] (v) Lyumjev (Eli Lily).[4, 11] Predicted monomeric lispro in humans has been adapted from pharmacokinetic modeling from our previous work in pigs.[8]

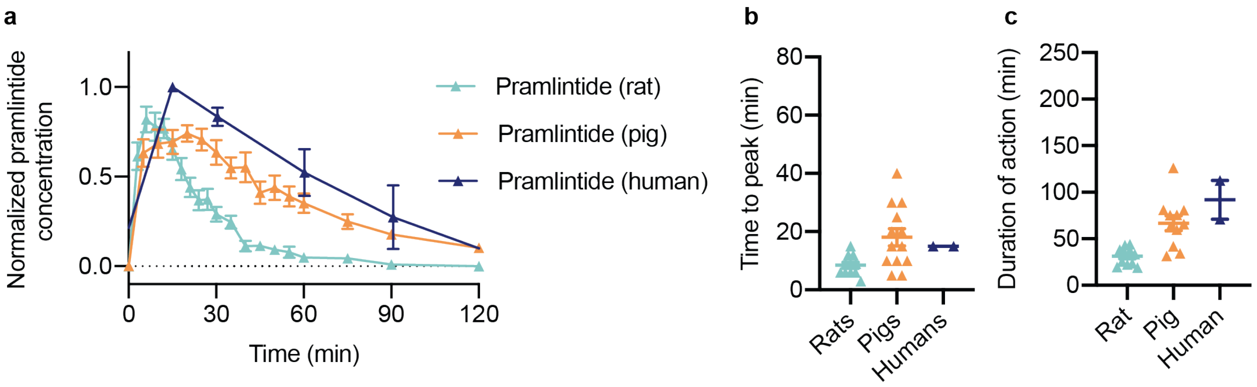


**Figure S9. Pramlintide pharmacokinetics for different species. a,** Normalized pharmacokinetics for pramlintide delivered in rats, pigs and humans. **b,** Time-to-peak for pramlintide delivered in rats, pigs and humans. **c,** Duration-of-action for pramlintide delivered in rats, pigs and humans. The conservation of the ultra-rapid absorbance kinetics from rats to pigs and the similar pramlintide kinetics between pigs and humans corroborates the model predicted kinetics for monomeric lispro in humans. Duration of action is defined here as peak width at 25% peak height ($time_{25\%\ down}$ - $time_{25\%\ up}$). Rat data is taken from this study. Pig data was adapted from previous work.(12) Human pramlintide data is adapted from two external studies.(13, 14)

**Table S1. MoNi copolymer excipient characterization.**

| Carrier Monomer | wt.% (Target) | wt.% by NMR (Exp) | Dopant Monomer | wt.% (Target) | wt.% by NMR (Exp) | $M_n$[a] (Da) | $M_w$[a] (Da) | Đ[a] |
|---|---|---|---|---|---|---|---|---|
| Acryloylmorpholine (Mo) | 77 | 74.5 [b] | N-isopropylacrylamide (Ni) | 23 | 25.5 [b] | 3200 | 3800 | 1.19 |

[a] Determined using Size Exclusion Chromatography calibrated using polyethylene glycol samples.

[b] Weight percentages difficult to determine due to overlapping spectra. Weight percentages estimated from post-precipitated NMR spectra by measuring the more resolved left half of the peak of N-isopropylacrylamide (δ= 4.0, 0.5 H), doubling it, and subtracting it from the unresolved peaks of Mo and Ni (δ= 3.2-4.2, 7H (Mo) 1H (Ni)).

### REFERENCES


1. J. Plank, A. Wutte, G. Brunner, A. Siebenhofer, B. Semlitsch, R. Sommer, S. Hirschberger, T. R. Pieber, A Direct Comparison of Insulin Aspart and Insulin Lispro in Patients With Type 1 Diabetes. *Diabetes Care* **25**, 2053 (2002).

2. R. J. Pettis, L. Hirsch, C. Kapitza, L. Nosek, U. Hövelmann, H.-J. Kurth, D. E. Sutter, N. G. Harvey, L. Heinemann, Microneedle-based intradermal versus subcutaneous administration of regular human insulin or insulin lispro: pharmacokinetics and postprandial glycemic excursions in patients with type 1 diabetes. *Diabetes Technol. Ther.* **13**, 443-450 (2011).

3. G. Andersen, G. Meiffren, D. Lamers, J. H. DeVries, A. Ranson, C. Seroussi, B. Alluis, M. Gaudier, O. Soula, T. Heise, Ultra-rapid BioChaperone Lispro improves postprandial blood glucose excursions vs insulin lispro in a 14-day crossover treatment study in people with type 1 diabetes. *Diabetes Obes. Metab.*, (2018).

4. H. Linnebjerg, Q. Zhang, E. LaBell, M. A. Dellva, D. E. Coutant, U. Hövelmann, L. Plum-Mörschel, T. Herbrand, J. Leohr, Pharmacokinetics and Glucodynamics of Ultra Rapid Lispro (URLi) versus Humalog® (Lispro) in Younger Adults and Elderly Patients with Type 1 Diabetes Mellitus: A Randomised Controlled Trial. *Clin. Pharmacokinet* **59**, 1589-1599 (2020).

5. L. Heinemann, T. Heise, L. C. Wahl, M. E. Trautmann, J. Ampudia, A. A. R. Starke, M. Berger, Prandial Glycaemia After a Carbohydrate-rich Meal in Type I Diabetic Patients: Using the Rapid Acting Insulin Analogue [Lys(B28), Pro(B29)] Human Insulin. *Diabetic Medicine* **13**, 625-629 (1996).

6. A. Lindholm, J. McEwen, A. P. Riis, Improved postprandial glycemic control with insulin aspart. A randomized double-blind cross-over trial in type 1 diabetes. *Diabetes Care* **22**, 801 (1999).

7. K. Rave, O. Klein, A. D. Frick, R. H. A. Becker, Advantage of Premeal-Injected Insulin Glulisine Compared With Regular Human Insulin in Subjects With Type 1 Diabetes. *Diabetes Care* **29**, 1812 (2006).

8. J. L. Mann, C. L. Maikawa, A. A. A. Smith, A. K. Grosskopf, S. W. Baker, G. A. Roth, C. M. Meis, E. C. Gale, C. S. Liong, S. Correa, D. Chan, L. M. Stapleton, A. C. Yu, B. Muir, S. Howard, A. Postma, E. A. Appel, An Ultra-fast Insulin Formulation Enabled by High Throughput Screening of Polymeric Excipients. *Sci. Transl. Med.* **12**, eaba6676 (2020).

9. M. Fath, T. Danne, T. Biester, L. Erichsen, O. Kordonouri, H. Haahr, Faster-acting insulin aspart provides faster onset and greater early exposure vs insulin aspart in children and adolescents with type 1 diabetes mellitus. *Pediatr. Diabetes* **18**, 903-910 (2017).

10. T. Heise, T. R. Pieber, T. Danne, L. Erichsen, H. Haahr, A Pooled Analysis of Clinical Pharmacology Trials Investigating the Pharmacokinetic and Pharmacodynamic Characteristics of Fast-Acting Insulin Aspart in Adults with Type 1 Diabetes. *Clin. Pharmacokinet.* **56**, 551-559 (2017).

11. M. Shiramoto, R. Nasu, T. Oura, M. Imori, K. Ohwaki, Ultra-Rapid Lispro results in accelerated insulin lispro absorption and faster early insulin action in comparison with Humalog(®) in Japanese patients with type 1 diabetes. *J. diabetes investigation* **11**, 672-680 (2020).

12. C. L. Maikawa, A. A. A. Smith, L. Zou, G. A. Roth, E. C. Gale, L. M. Stapleton, S. W. Baker, J. L. Mann, A. C. Yu, S. Correa, A. K. Grosskopf, C. S. Liong, C. M. Meis, D. Chan, M. D. Troxell, D. M. Maahs, B. A. Buckingham, M. J. Webber, E. A. Appel, A co-formulation of supramolecularly stabilized insulin and pramlintide enhances mealtime glucagon suppression in diabetic pigs. *Nature Biomedical Engineering* **4**, 507-517 (2020).

13. O. G. Kolterman, S. Schwartz, C. Corder, B. Levy, L. Klaff, J. Peterson, A. Gottlieb, Effect of 14 days' subcutaneous administration of the human amylin analogue, pramlintide (AC137), on an intravenous insulin challenge and response to a standard liquid meal in patients with IDDM. *Diabetologia* **39**, 492-499 (1996).

14. M. C. Riddle, K. C. J. Yuen, T. W. de Bruin, K. Herrmann, J. Xu, P. Öhman, O. G. Kolterman, Fixed ratio dosing of pramlintide with regular insulin before a standard meal in patients with type 1 diabetes. *Diabetes Obes. Metab.* **17**, 904-907 (2015).
